# ABCnet : Self-Attention based Atom, Bond Message Passing Network for Predicting Drug-Target Interaction

**DOI:** 10.1101/2021.12.27.474154

**Authors:** Segyu Lee, Junil Bang, Sungeun Hong, Woojung Jang

## Abstract

Drug-target interaction (DTI) is a methodology for predicting the binding affinity between a compound and a target protein, and a key technology in the derivation of candidate substances in drug discovery. As DTI experiments have progressed for a long time, a substantial volume of chemical, biomedical, and pharmaceutical data have accumulated. This accumulation of data has occurred contemporaneously with the advent of the field of big data, and data-based machine learning methods could significantly reduce the time and cost of drug development. In particular, the deep learning method shows potential when applied to the fields of vision and speech recognition, and studies to apply deep learning to various other fields have emerged. Research applying deep learning is underway in drug development, and among various deep learning models, a graph-based model that can effectively learn molecular structures has received more attention as the SOTA in experimental results were achieved. Our study focused on molecular structure information among graph-based models in message passing neural networks. In this paper, we propose a self-attention-based bond and atom message passing neural network which predicts DTI by extracting molecular features through a graph model using an attention mechanism. Model validation experiments were performed after defining binding affinity as a regression and classification problem: binary classification to predict the presence or absence of binding to the drug-target, and regression to predict binding affinity to the drug-target. Classification was performed with BindingDB, and regression was performed with the DAVIS dataset. In the classification problem, ABCnet showed higher performance than MPNN, as it does in the existing study, and in regression, the potential of ABCnet was checked compared to that of SOTA. According to experiments, for Binary classification ABCnet have an average performance improvement of 1% for higher performance on DTI task than other MPNN and for regresssion ABCnet have CI with an average 0.01 to 0.02 performance degradation compared to SOTA. https://www.overleaf.com/project/618a05533676801d8f68ccf6

## Introduction

The study of the interaction between compounds and proteins plays an important role in the development of a wide range of drugs, as it can reveal the extent of the therapeutical benefit to patients of the activation, inhibition, or conformational changes of functional proteins. Drug-target affinity (DTA) predicts the interaction, specifically the binding affinity, between a drug candidate compound and a target protein, and it is possible to derive drug candidates which inhibit the activity of target proteins corresponding to diseases [4]. Developing a new drug takes an average of 5.5 years, and recent research shows that the average cost of developing an FDA-approved drug has been $1.3 billion over the past ten years, and such development will become even more costly after the COVID-19 pandemic [28]. Accordingly, many large pharmaceutical companies are conducting research in collaboration with artificial intelligence ventures to reduce costs and time by introducing artificial intelligence technology to new drug development.

There are two methods for DTA prediction: in vitro and in silico. Although in vitro is accurate, it is expensive and time-consuming, while in silico saves time and money by performing virtual screening using a computer prior to experimental verification. There are two types of in silico prediction methods of drug-target interaction: molecular docking and machine learning. Molecular docking utilizes 3D simulation to identify potential binding sites within the structures of prospective drug molecules and targets them with high accuracy and visual interpretation. However, in silico methodology cannot be applied when the three-dimensional structure of a protein is not known, and large-scale simulations using this method require a lot of time.

Machine learning approaches are attracting attention because they can scan a large number of candidates in a short period of time [2].An immense volume of chemical and biomedical data have accumulated over decades of experiments, prompting the emergence of a data-based research methodology, and recently, a machine learning method that increases performance according to the quantity and quality of data is attracting attention [21]. The deep learning method has achieved overwhelming success in fields such as computer vision, natural language processing, and speech recognition in particular [10]. As the applicability of deep learning has been proven in various fields, research on its application to drug discovery has also emerged, first introduced in the fields of molecular property prediction and DTI [1]. Recently, research on the latest technology beyond the introduction stage is also being conducted, including cases where GCN [15] and transformer [12] methodologies are applied to DTA and molecular property prediction. Recent studies have been conducted using deep learning methods to find drug candidates for the SARS-CoV virus [29] [3]. Since deep learning can find drugs which can interact with and bind to disease-causing target proteins in a relatively short time and at low cost, deep learning in the discovery of candidate substances during drug development is attracting attention.

DTA is an analysis which predicts the binding affinity of a compound and a target by receiving the sequence data of a compound and a target protein, and can be applied to various drug discovery processes. To explain the relationship between DTA input and output, compound data, a component of the input data, is commonly represented with a data expression method called SMILES, a format which lists compound element symbols and combinations as strings. The target protein is expressed as a one-dimensional sequence, in which its amino acids are listed in the form of a string. From these two strings, the DTA model learns and predicts whether or not the compound and target protein will bind with each other, and to what extent.

The architecture of the deep learning-based DTA model consists of three major components. The first is the drug encoder, which converts the information about atoms and bonds expressed as strings in SMILES into a data form that can be learned with RDkit from a generated one-hot encoding matrix. In other words, it is a step in generating expression embeddings which can better contain structural and physical characteristics of compounds by using the simple compound expression data format SMILES.

The second is the target encoder, which converts the amino acid sequence of a protein into a one-hot vector for each amino acid expressed in 26 amino acid combinations, then converts the sequence into a vector expression. That is, it performs the expression embedding of the protein. The third is a decoder which fuses the drug and protein feature embedding results extracted by the drug and target encoders into a latent vector. Several deep learning techniques can be used in the subsequent process. If the prediction target is regression, the model is configured to yield an affinity score if it is classified and whether it is combined.

Our study aimed to derive performance beyond the SOTA by focusing on improving the drug encoder during the DTA stage. Various efforts are underway to extract drug characteristics. Among them, research using message passing neural networks (MPNNs) have been active for several years. Gilmer [8] first introduced MPNNs in the effective extraction of drug characteristics, Tang [24] implemented self-attention in addition to MPNN, and Huang [11] performed a study to improve MPNN. MPNN creates the message passing function through the Dense layer before creating the initial value, that is, the 0th hidden state for the atom. And without calculating the hidden state of the bond, based on only the atom *v*, only the information about the neighboring Atom *u* and the neighboring Bond *v* − *u* is transferred to create an atom message. This method makes it difficult to properly reflect the structural information of the compound due to the simplification of Atom initial value setting, and since the bond also uses only features, the structural information about the bond is ignored. Huang’s study added a hidden state calculation for the bond to improve the bond constraint, but it did not overcome the limit for the atom.

In our study, we did not set the initial value of the atom hidden state as a simple dense layer, but used the neighboring atom feature information to set the initial value using the dense layer, message passing, and update function. The second point of focus, the Bond feature, is not the method that uses the existing raw data as it is, but the method of setting the initial value of the atom hidden state is also applied to the bond. In addition, referring to the study using self-attention, self-attention was used in Atom, Bond, and Concat Message to improve the part where the relationship between data that is too far away, which may occur in sequence data, is not well reflected.

We propose ABCNet as Figure 1. First, the Drug Encoder is composed of an Atom initial message block, a Bond initial message block, and a Concat message block that concatenates the Atom and Bond initial hidden state, transmits neighboring Atom and Bond information, and generates an Atom message. This is called ABCnet: Atom, Bond, Concat message passing network.The second is a Target Encoder, which uses a Simple Convolutional Neural Network as a protein model to extract protein features.The second is a Target Encoder, which uses a Simple Convolutional Neural Network as a protein model to extract protein features. The third is a decoder, which predicts a drug-target interaction score through a fully connected layer by using a latent vector obtained by concatenating drug and protein features extracted from the drug encoder and the target encoder as input data.

**Figure 1.**
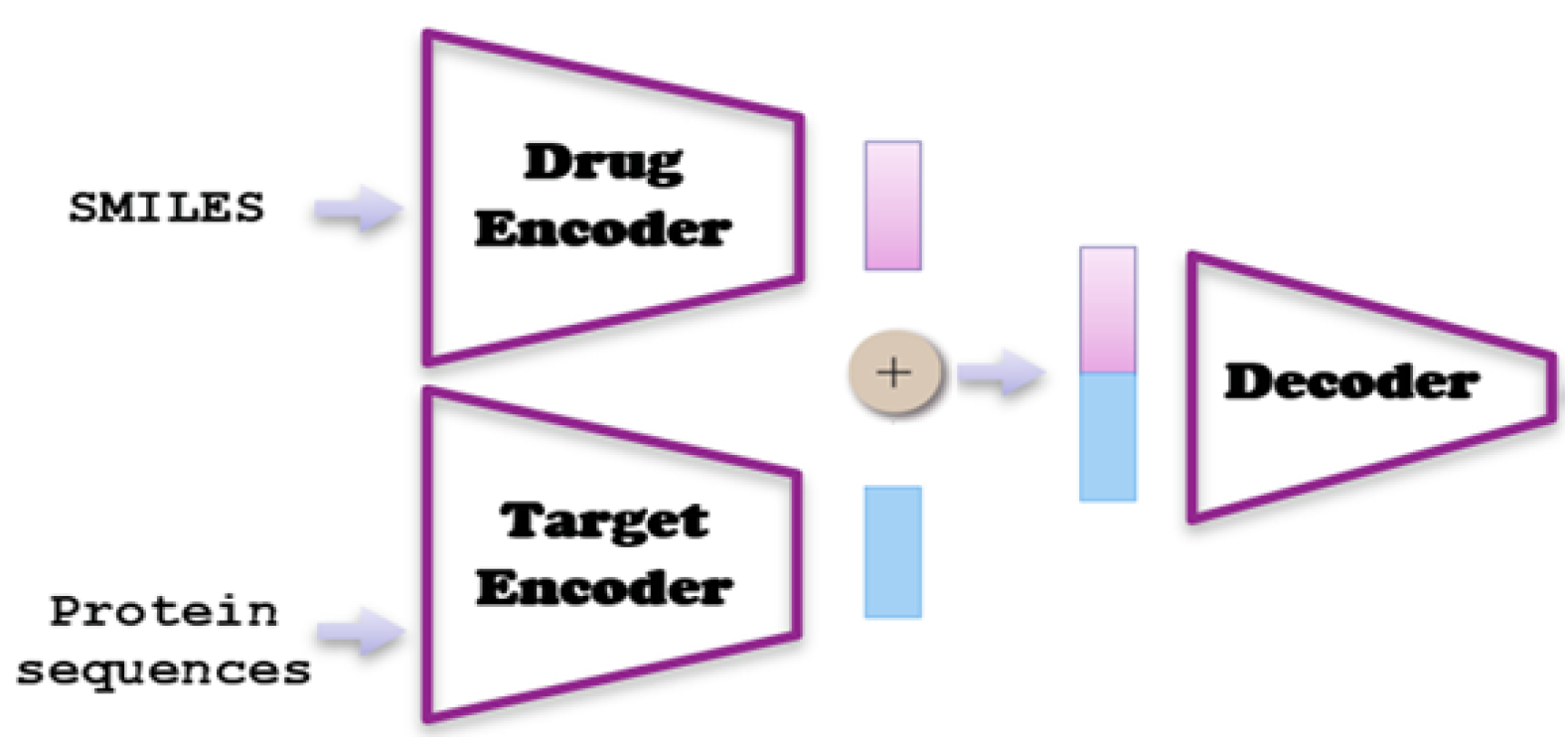
Example of a standard floating figure. **A-F**, This figure is wrapped into the standard floating environment.

**Figure 2.**
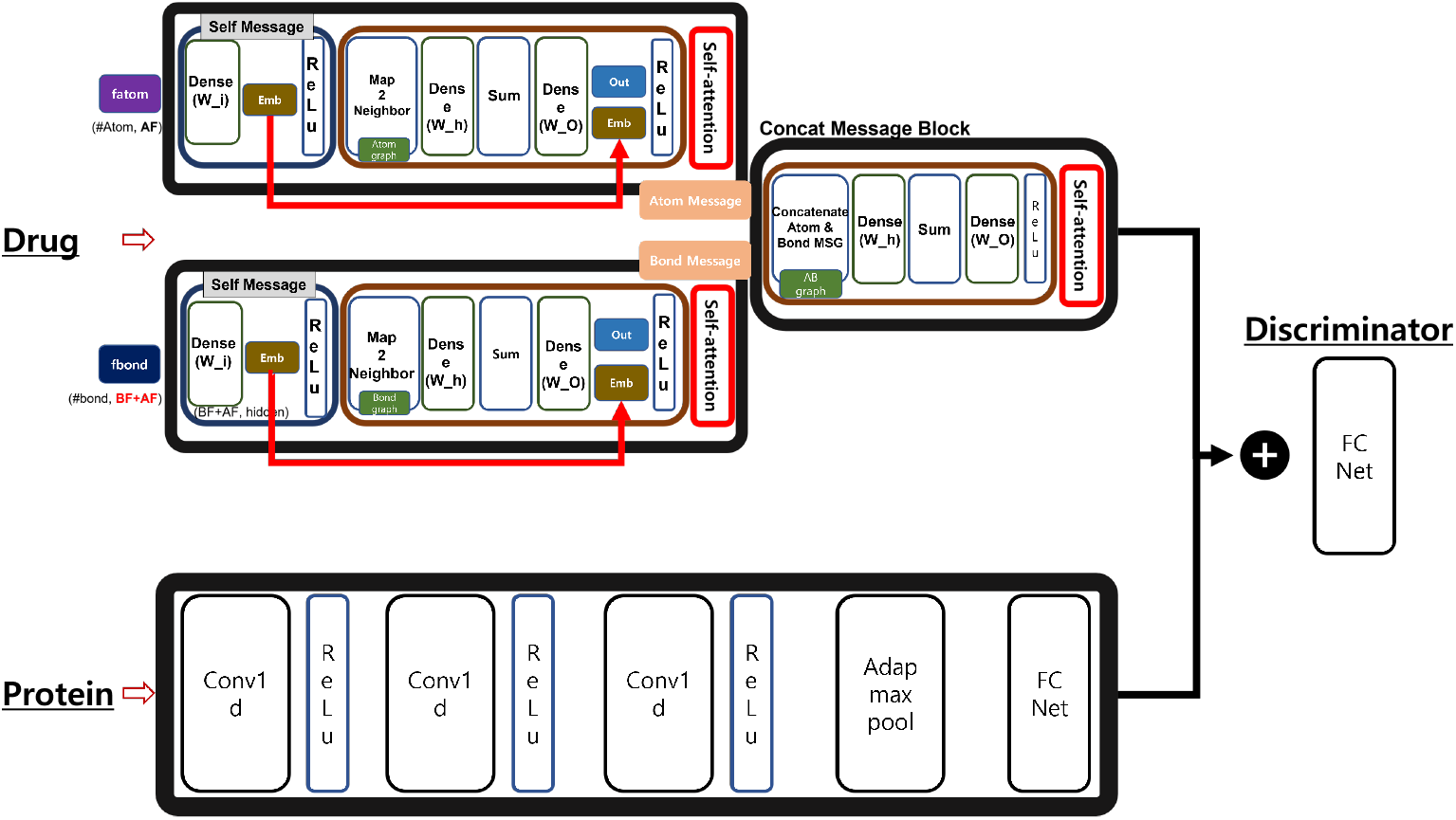
ABCNet overview.

**Figure 3.**
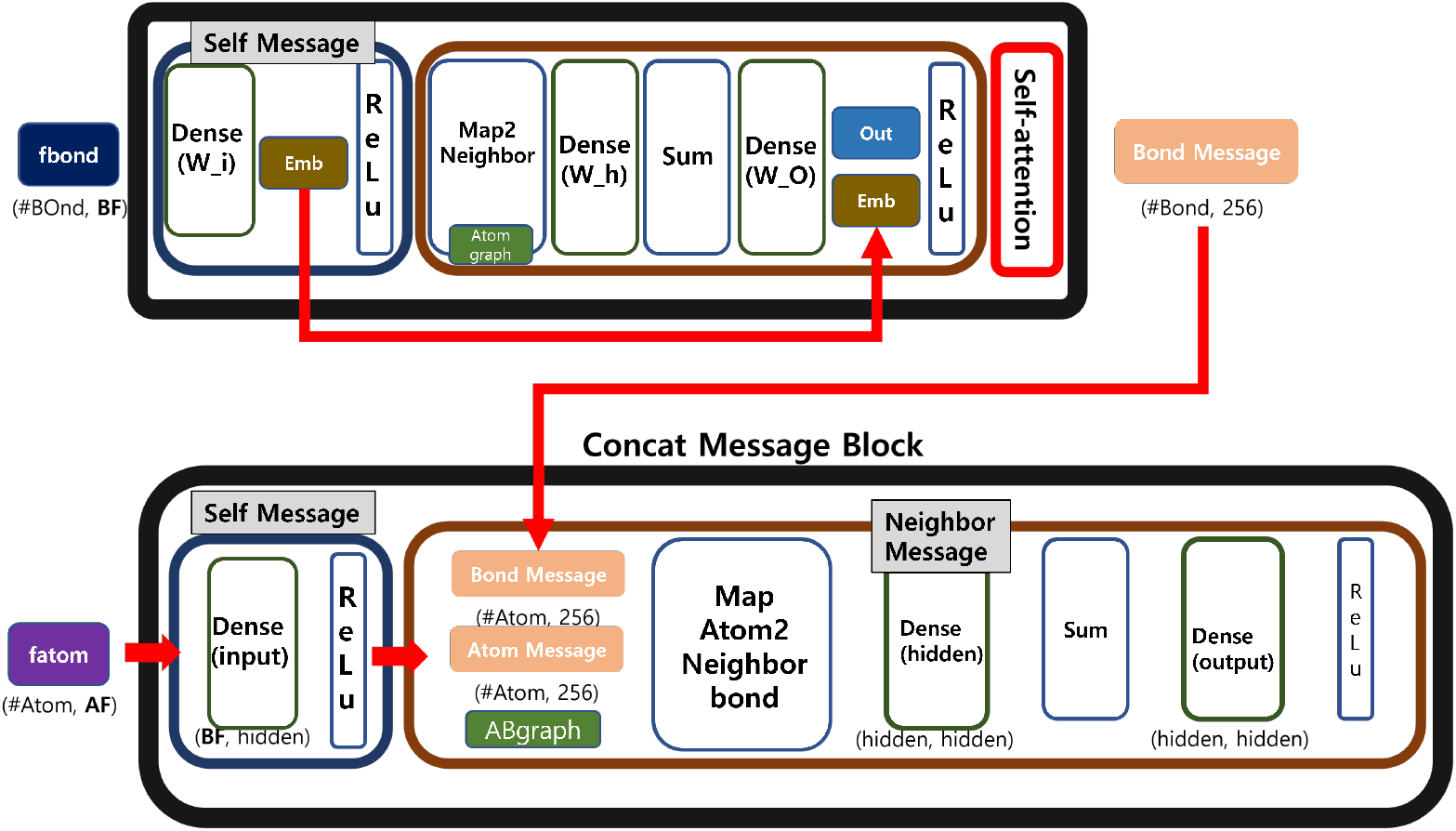
BCNet overview.

**Figure 4.**
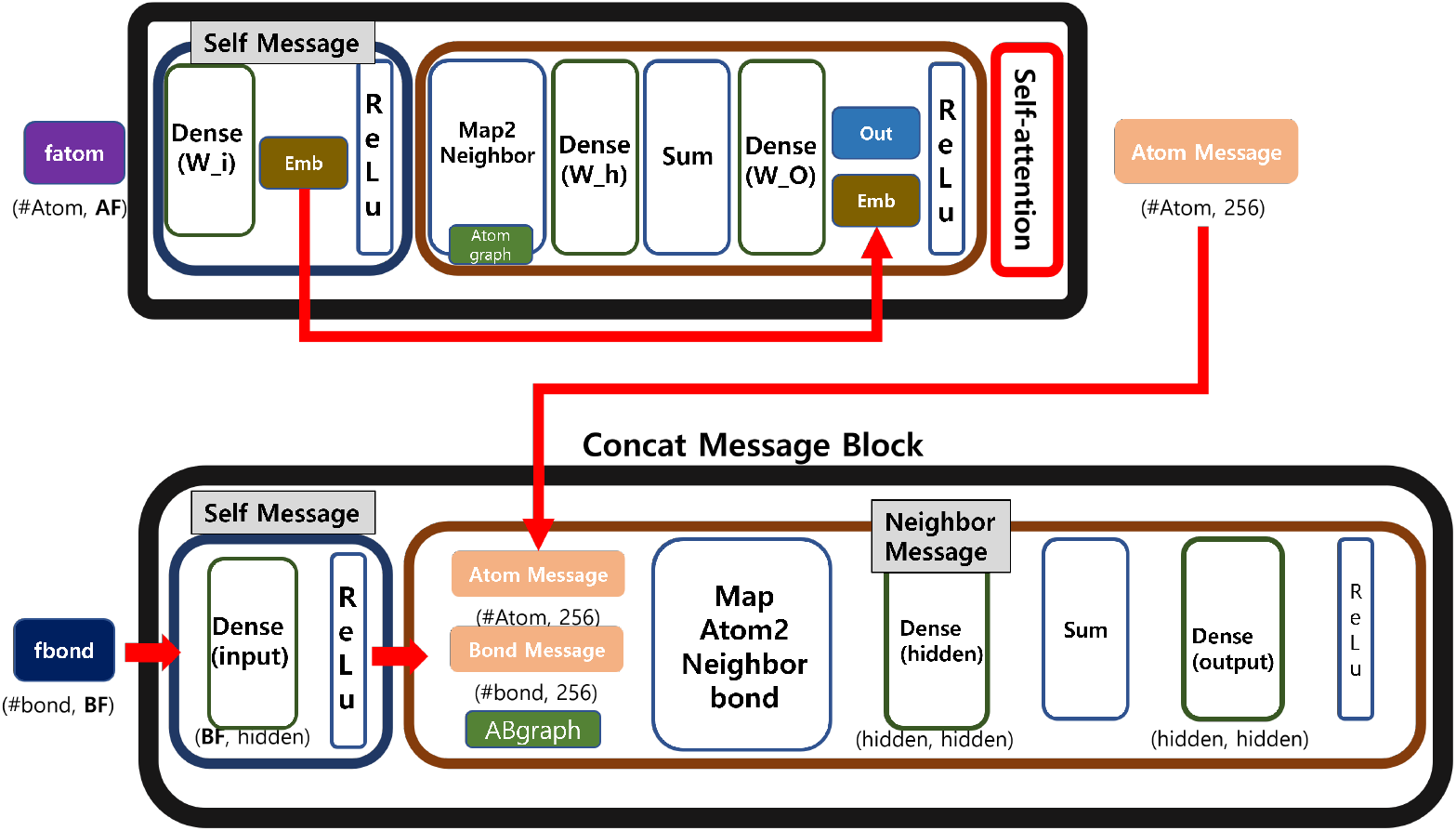
ACNet : Atom, Bond, Concat Message Passing Neural Network.

**Figure 5.**
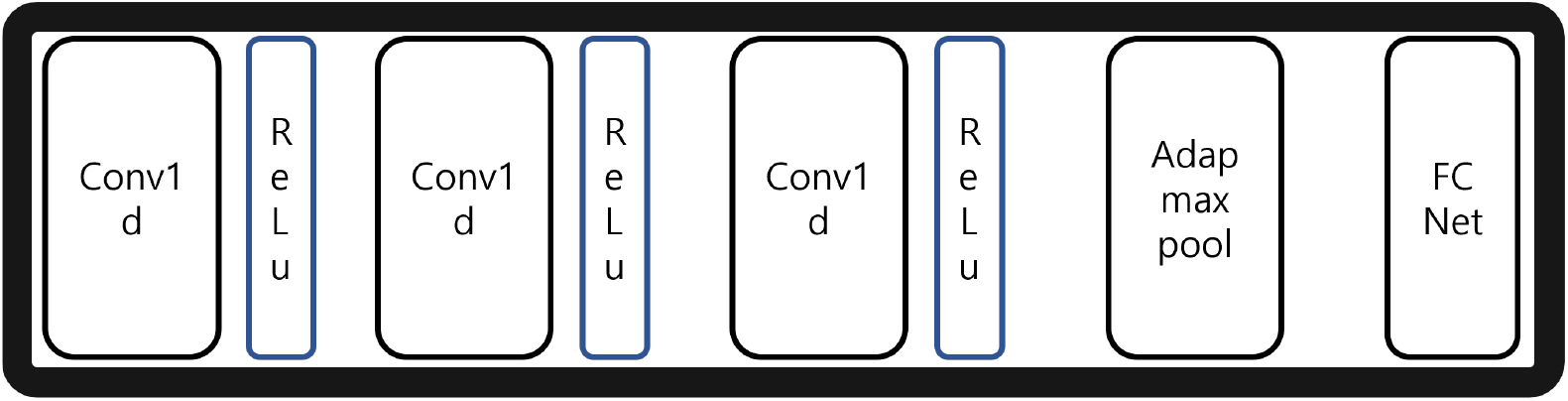
Protein Model overview.

To verify our proposed ABCNet, two experiments were performed: Binary Classification, which predicts DTI, and Regression, which predicts the binding score.BindingDB [17] and DAVIS [5] benchmarks were used for each experiment, and accuracy and AUPR were used as performance evaluation indicators, and MSE and AUPR were used. For accurate performance comparison, SOTA algorithms were performed in the same environment and the results were compared and analyzed. As a result of performance evaluation with SOTA, there was a 1-2% improvement in classification and 0.01 0.02 lower performance in regression CI.

The purpose of the Binary Classification experiment is to compare existing MPNN models to find out whether the drug encoder of the proposed model extracts drug properties well. In order to achieve the objective, the structure of the rest of the models except for the drug encoder and the hyperparameter values were matched to compare the drug encoder to find the optimal drug encoder. Since this result performance is all the same except for the drug encoder, the DTI performance result depends entirely on the drug encoder. Through regression analysis with ABCnet derived in this way, the performance was compared with State-of-the-art (SOTA) of DTI.

### Related works

As studies of predicting drug-target protein interaction (DTI) using artificial intelligence have been actively conducted, these two studies have received a lot of attention. The first study, DeepDTA, is the first study to predict the interaction using deep learning by binary classification. This study proposed a deep learning-based 2-way CNN model that uses only the sequence information of the target and drug to predict the interaction binding affinity, and utilized the Davis and KIBA dataset. Through this study, it can be seen that the door to new drug development research using artificial intelligence has been opened, and it is being used as a model that serves as a reference point for other studies. The second study, GraphDTA, is a model that graphs drugs and predicts drug-target affinity using a graph neural network. The graph was used so that the data in the original graph form could be used in a way similar to the natural state, rather than using a simple sequence. This study also used the same data as DeepDTA. Through this study, it was verified that graphs are a method that can be used for learning by better representing drug data. The detailed application method of the graph-based deep learning model used in this study and related references are as follows.

Graph Convolution Neural Network (GCN) refers to a model in which the convolution operation applied to the existing image pixel values is applied to the graph. There are two main types of GCN: Spectral-based GCN and Spatial-based GCN. Spectral-based GCN is a method of convolution by introducing a filter as in signal processing operation, and Spatial-based GCN is a method of convolution using node features. More specifically, spectral-based GCN performs eigen decomposition of Laplacian matrix on the graph as if filter operation is applied in existing signal processing.

The Laplacian matrix can understand the degree and connection relationship between each node, that is, how the nodes in the graph are connected to each other. By decomposing this Laplacian matrix by eigenvalues, it becomes possible to express signal transmission between nodes. However, since such a spectral-based GCN method has to input all nodes of the graph at the same time, it can consume a lot of time and cost when a large-sized scrap is input. In order to improve this limitation, Spatial-based GCN can save a lot of time cost compared to Spectral-based GCN by performing convolution only with neighboring nodes. By decomposing the Laplacian matrix by eigenvalues, it is possible to express signal transmission between nodes. However, since such a spectral-based GCN method has to input all nodes of the graph at the same time, it can consume a lot of time and cost when a large-sized scrap is input. In order to improve this limitation, Spatial-based GCN can save a lot of time cost compared to Spectral-based GCN by performing convolution only with neighboring nodes.

The GCN used in this paper does not input all nodes in the graph at the same time in the spectral-based GCN, but introduces the Chebyshev polynomial and limits only the K-th neighbor nodes to greatly reduce the time complexity. When there is a graph Graph(N, E) composed of N nodes and E edges, by embedding each node in d dimension, an adjacency matrix expressing the n x d dimension input value and the connection relationship between each node can be obtained. For *Graph*(*N, E*), it is expressed as Input *X* : *n* x *d*, and the adjacency matrix *A* : *n* x *n*. (Where *N* is Node(Atom), *E* is Edge(Bond), *n* is the number of all Node, *d* is the feature dimension for Node and *A* is the adjacency matrix for the graph). In this paper, using these input x and adjacency matrix A as input data, a forward pass was performed through a filter calculated by applying the Chebyshev polynomial and two hidden layer GCNs. Through this hidden state, node-level prediction is made. Through node-level prediction, we can classify semi-supervised nodes for nodes we do not know [14].

Message Passing is a necessary process in most Graph Neural Networks. Among the methodologies for learning graphs, the Spatial Convolutional Network borrows the idea of a Convolutional Neural Network, and CNN combines the surrounding pixels using a convolution filter when determining the value of the central pixel. In the Spatial Convolutional Network, it works by merging the features of neighboring nodes instead of neighboring pixels. In addition, the Spectral Convolutional Network, designed based on the graph signal processing theory, also performs the matrix multiplication of the adjacency matrix and the feature matrix, which are data representations of the graph. Most Graph Convolutional Networks (GCNs) operate in this way, and research has been conducted on how to transmit information to share and update node information.

The function corresponding to how information is transmitted is the message-passing function, and Gilmer Justin conducted a study to apply GNN to Quantum Chemistry using the following three functions. In this study, we describe an MPNN that has a message passing phase and a readout phase in forward propagation, and operates on an undirected graph G for simplicity. The message passing phase is shown below. In the equation, node features are *x*_*v*_, edge features are *e*_*vw*_, message function is *M*_*t*_, vertex update function is *U*_*t*_, hidden states 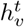 at each node in the graph, the updated message is 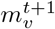, and *N* (*v*) is the neighbors of v in graph *G*.

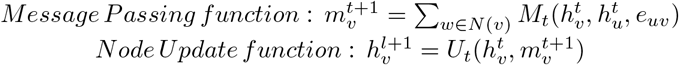

In the readout phase, the message function is *M*_*t*_, the vertex update function is *U*_*t*_, and the readout function *R* is used to calculate the feature vector for the entire graph and is expressed by the following equation.

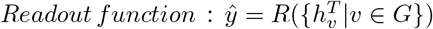

The first step is to deliver the neighbor message to its own node. The second step is to update the own node message through the message received from the neighbor. The feature is that messages from neighboring nodes are reflected by using an adjacency matrix. The parameter that determines how many neighboring nodes are viewed is called hop or depth, and according to this size, all the features of the subgraph of the entire graph are transmitted to compose one node information [8].

As mentioned in the MPNN section above, MPNN studies are very important for learning graph-structured data. This is because the embedding performance, which is a new expression created as a by-product of learning, varies greatly depending on how the MPNN method is designed. In Withnall’s study, there is a difference in the method of creating edge features when constructing neighboring messages, unlike the existing MPNN method. The difference is that when constructing a neighboring message, when creating an edge feature, the edge feature is embedded by concatenating the edge’s own feature and two node features connected to the edge. By adding two node features connected to the edge in this way, the edge feature information is enriched [27].

Kearnes [13], Huang [11] conducted a study that modified the edge feature learning method in various ways. In this study, edge features were learned by setting hidden states for edges separately. When updating an edge message, not only the immediately preceding neighbor (1-hop) edge message is obtained as in the existing MPNN, but an edge message that is as far away as the depth (n-hop) is obtained, and more neighbors are considered in the configuration of the edge message, and the amount of information around to expand the expressive power of embedding learned with MPNN.

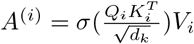

Input data includes *d*_*k*_ dimension Query *Q*_*i*_, Key *K*_*i*_, and Value *V*_*i*_. Calculates the dot product for input data, query and key. This is to find the similarity between each Query *Q*_*i*_ and Key *K*_*i*_ through dot-product.Divide the calculated value by 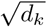. By scaling with 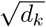, it is to prevent in advance the situation in which there is little change in the derivative value of the Softmax function as the dot-product value increases. Using this calculated value, the weight for Value *V*_*i*_ is obtained through Softmax. If this weight, that is, attention score, is multiplied by Value *V*_*i*_, the value similar to Query *Q*_*i*_ has a higher value. In other words, it is possible to express the attention mechanism that the more similar the value, that is, the more important the information, the more focused on that information.

Attention was first introduced in Vaswani’s study.Previous sequence data were trained with a Recurrent Neural Network. RNN, which is a method of learning by remembering data order, can reflect the order of sequence data or a neighbor relationship, so it has been used for many sequence data. However, the RNN has to receive a sequence unit input, and when the length increases, the gradient is lost due to continuous matrix operation, so there is a problem that the relationship with distant data is not properly reflected. Attention is a way to solve this problem. In a way that does not only consider the distance of a sequence, but rather considers information about the relationship between words, high correlation can be shown even if they are far apart. Attention, unlike RNNs, does not learn sequentially, so it also has a computational advantage [25]. In particular, attention has achieved high performance in the field of natural language processing, and the famous BERT [6] and GPT [22] are also attention-based algorithms.

Shin applied a Transformer that includes attention to DTA.The information included in SMILES, which is a string expression of molecules, was transformed so that it can be used as an input to the Transformer.This study proved that the Transformer technique can be applied to the embedding of important compounds in the bio field [23].

Similarly, Maziarka added a total of three features of Molecule distance, adjacency, and self-attention to learning when applying the Transformer to Molecule property prediction. [18]. These studies have shown that attention can improve SOTA in predicting the properties of compounds or binding affinity with the target. Additionally, there is a study by Tang that applied self-attention to a message passing neural network (MPNN) to predict chemical properties. Basic MPNN was applied to Molecule, and self-attention was applied to the feature information matrix for each atom that came out through MPNN [24].

In this study, using this self-attention method, we studied a model that preserves atom and bond information through self-referential calculation formula in feature extraction and reduces information loss after calculating adjacent nodes. By scoring each atom and bond in self-attention, it is designed to identify factors that significantly contribute to or degrade the interaction. In the next, experiments to add methods to the protein model were adopted as the default algorithm for upgrading each data to a descriptive artificial intelligence model. In this paper, we propose a method for binding affinity prediction of protein and molecules using self-attention graph based Drug Target Interaction 2-way, end-to-end Model. Here, 2-way refers to a model that extracts drug and protein features, respectively. After passing through a model that predicts drug-target binding by concatenating two latent vectors containing characteristic information of drugs and proteins that have been obtained through this, binding presence (Binary) or binding value (Regression) is output.When delivering a message using self-attention, attention is given to deliver a message with high relevance to drug-target combination. As a result, we proposed the ABC structure, which is the drug feature extraction MPNN structure that can best predict DTI.

### ABCNet

ABCnet (Self-attetion based Atom, Bond, Concat message passing network), unlike the previous MPNN, is divided into two parts: creating, delivering, and updating messages.The first is Atom, which creates, delivers, and updates Messages about Atom. The second part is Bond, which creates, delivers, and updates messages about Bond.The third concatenate network proceeds based on the atom message and the bond message. The message is created separately by dividing the single message generation method created in the previous MPNN into Atom and Bond. In this way, Atom can grasp Molecule structural information based on Atom, and Bond can grasp Molecule structural information based on Bond. Concatenate the found Atom and Bond standard Molecule structural information (Message) and combine the previously calculated Atom and Bond messages based on the connection relationship between Atom and Bond within the Molecule in the Concat message block. Structural Molecule features are extracted with a Drug Encoder by applying self-attention to the calculated results.

ABCnet is a model that predicts binding affinity using the fully connected layer in the decoder by concatenating the drug and protein feature vectors calculated using CNN in the drug encoder and the target encoder, respectively.

### Drug Model

The drug model uses SMILES as input data, but it does not encode SMILES. The Atom, Bond Feature matrix, which has extracted features for Atom and Bond, is used as input data. Atom and Bond features were extracted using the Python library of RDkit [16]. Gilmer’s research [8] and Tang’s research [24] were referenced for specific features to be extracted from atom and bond.

We tried to extract features that reflect Molecule’s properties and features that are highly related to drug-target binding. Formal Charge, Degree, Chirality, Aromaticity, Bond Stereo, Ring, which contain the chemical and structural characteristics of Molecule, including atom and bond types that must be entered, were selected.

Furthermore, the hybridization type, that is, the orbital shape in which electrons move, was used as a feature in order to consider the electrical characteristics and shape that can have an important effect on drug-target binding.

### BCNet: Bond, Concatenate Message Passing Neural Network

Previous MPNN studies used only Atom Messages without Bond Messages. BCnet adds the idea of bond-oriented message creation and delivery. Since Molecule is composed of Atom and Bond, Bond was considered as an important factor in drug-target binding as much as Atom. So, the Bond message passing part that creates and delivers the Bond message is separated and created. Just as Atom is connected to neighboring Atom, Bond is also connected to neighboring Bond, so the Bond message was created in the same way as creating an Atom message. Just as Atom is connected to neighboring Atom, Bond is also connected to neighboring Bond, so the Bond message was created in the same way as creating an Atom message. Bond message made based on the Molecule connection relationship not only contains simple bond features, but also includes feature information about surrounding bonds, so it reflects not only simple bond feature information but also structural information on the bond graph. Bond message replaced the raw bond feature used in previous MPNN research, and the bond message reflecting bond feature and graph structural information can express Molecule better than before. The MPNN operates in the following way, through the *t*th hidden state for its own Bond *v* and the *t*th hidden state for its neighbor Bond *w* through the *t* + 1th hidden state for its own *v* create a message.

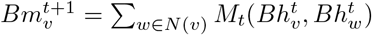

The *t* + 1th message for the renewed self Bond *v* and the previous *t*th Bond hidden state for the self *v* bond hidden state for the *t* + 1th own Bond *v* update.

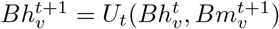

After that, the hidden state for Atom *v, w* and *v*−*w* Bond hidden state are input to create a Message.

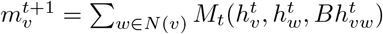

A new hidden state for the t+1th atom v is created through the message about atom v and the previous t-th atom v hidden. The feature vector is calculated through the hidden state of the atom v that has been calculated in this way.

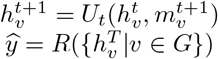

### ACNet : Atom, Concat Message Passing Neural Network

ACNet performs the same process as the existing MPNN research, but by making the layer that creates and delivers the atom message deep, it can learn a higher-order expression than the existing method. When setting the initial value of the hidden state, instead of using a dense layer, various information can be learned because the layer is deep to include the structural information of the graph about the atom. The procedure and the Bond MPNN method introduced earlier are the same, but the target is Atom.

1. Atom Message Block atom message : Creates an atom-based message by receiving neighboring atom features as much as Depth. Then, the atom message of *t* + 1 is updated with the atom message and hidden state. Finally, it creates its own atom message with the neighbor atom message as much as depth and its own atom message, and updates the message to be sent as the atom message.

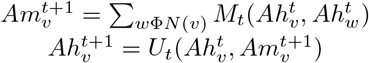
2. Bond Message Block bond message : A bond-based message is created by receiving neighboring bond features as much as depth. Then, the bond message *t* + 1 state is updated with the bond message and the hidden state. Finally, it creates its own bond message with the neighbor bond message and its own bond message as much as the depth, and updates the message to be sent as an atom message.

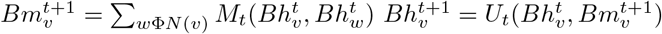
3. Concat Message Block concat message : An atom-based message is generated by receiving neighboring atom messages and bond messages as atom messages and bond messages. And calculate the feature vector with the hidden state of atom *v*.

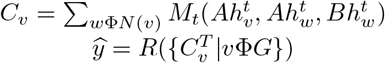

In our study, when generating a message, an atom-based message was created by receiving the neighboring atom features equal to the depth and the bond feature just before it. In the previous MPNN algorithm, since only the immediately preceding neighboring bond feature was referenced, there was a disadvantage in that the information of the bond message was limited, and we changed the model to consider both Atom and Bond message passing in message passing. In this process, since neighbor information is delivered as much as depth to each atom and bond message configuration, the features of the entire molecule can be delivered more abundantly.

At the end of MPNN, a vector for each atom is calculated through a readout function, and self-attention was added to this part to obtain two effects. First, self-attention was applied to the last vector of the Concat message block, in order to obtain an interpretable vector to find the atom of the drug-target binding part. Second, self-attention was applied to the last generated vector of atom message block and bond message block, for effective learning by assigning weights according to the degree to which substructures for each atom and bond contribute to drug-target binding.

ABCnet is similar to the structure of the existing DTI model. The first input is the drug SMILES data, and the second input is a one-dimensional protein sequence in which amino acids are expressed. As each input, we embed features in the Drug Model and Protein Model. In particular, in the Drug Model, Atom-based GCN and Edge-based GCN are calculated so that the graph data structure can be learned better. Here, Atom-based GCN is defined as ACnet, and Edge-based GCN component is defined as BCnet. Attention is added to each result so that importance can be considered. In order to predict the interaction, we fused the two results that went through attention and used a fully-connected layer. In addition, by adding attention at the end, it was possible to determine which drug atom and which protein amino acid is related in DTI. In other words, the design intention is to express the correlation between each element of Drug and Protein in the DTI numerically, and to enable an analytical model as a result. Consequently, we build Total model, which is Two-ways End-to-End Neural Network for predicting DTI.

### Protein Model

Protein is a sequence of several amino acids, which can be expressed in the form of one-dimensional amino acid sequence data. Sequence data can be expressed as a matrix in the form of one-hot encoding, and there are a total of 26 amino acids (include unknown). When expressing methionine, ‘M’ can be expressed as a vector having a size of (26, 1). If the protein sequence has 1,200 amino acids, it can be expressed as a size of (26, 1200). Similarly, protein sequences can have different lengths, and the length can be adjusted depending on whether the model wants a fixed length or not. Using a one-hot encoding matrix as input data, it passes through a one-dimensional convolutional neural network to embed a protein sequence. Since our study focused on the modification of the drug model, the protein model consisted of a simple extraction section.

### Experiment

Two problem situations were assumed for the performance evaluation of our model. Binary classification, where the label that predicts whether the drug-target is bound or not, can be expressed as 0 and 1, and regression, which predicts the IC50, which is the drug-target interaction score.

In Binary Classification, various MPNN models introduced and described above (previously MPNN, BCnet, ACnet, ABCnet) were all tested to find the MPNN model with the best performance. The model with the best performance was used for the regression problem experiment, and it was analyzed and compared with the DTI SOTA model. When comparing various MPNN models in binary, ABCNet, which is the best model, achieved more than 96.6% performance, and in the regression problem, it was compared with ABCnet and SOTA models.

### Dataset

In order to predict the DTI, experiments were conducted to predict the drug-target binding (binary) and drug-target binding score (regression). The experimental dataset for each of Binary and Regression is shown in Table 3 below.

**Table 1.** Atom Feature Table. This table lists the characteristics used to train the model.

| Attribute | Description | Dimension |
| --- | --- | --- |
| Atom type | Well-Known and Unknown chemical elements | 23 |
| Hybridization type | S, SP, SP2, SP3, SP3D, SP3D2 | 6 |
| Formal Charge | Charge assigned to an atom | 5 |
| Degree | Number of heavy atom neighbors | 6 |
| Chirality label | R, S, Unspecified and unrecognized type of chirality | 4 |
| Aromaticity | aromatic atom or not | 2 |

**Table 2.** Bond Feature Table. This table shows the Bond characteristics used for model training.

| Attribute | Description | Dimension |
| --- | --- | --- |
| Bond type | Single, Double, Triple, Aromatic | 4 |
| Bond Stereo | Bond stereo-chemistry type | 6 |
| Ring | Whether the bond is in a ring | 2 |

**Table 3.** This table shows the Bond characteristics used for model training.

| Problem | Dataset | SMILES | Protein Seq | Label |
| --- | --- | --- | --- | --- |
| Binary | BindingDB | O=Cc1ccc(O)c(OC)c1 | MEICAGIVEASV | 0 or 1 |
| Regression | DAVIS | O=Cc1ccc(O)c(OC)c1 | MEICAGIVEASV | Kd |

Binary Classification was performed with BindingDB Dataset [16]. BindingDB is a public, web-accessible database of measured binding affinities, focusing chiefly on the interactions of protein considered to be drug-targets with small, drug-like molecules. Interaction indicators of BindingDB include Ki and IC50. The experiment was carried out by selecting the IC50 with the largest number of data as the label. IC50 is an abbreviation of The inhibitory concentration of drug that causes 50% of the maximum inhibition, and represents an active action on drug-target binding. It refers to the concentration of the drug when the degree of binding between the drug and the target is suppressed to 50%. Therefore, a small IC50 value means that a drug has a high binding affinity since it reduces the degree of binding to a target by 50% even at a low concentration. The IC50 value of BindingDB is an integer, and there are several numbers. In order to proceed with model learning with binary classification, it is necessary to determine whether the label is 0 or 1, that is, whether it is not combined or whether it is combined. For the thresholds of positive and negative labels for binary classification, refer to [26] and [19] papers. According to these papers, Label is positive if its IC50 is less than 100nm, negative if IC50 greater than 10,000nm.

Regression was performed with DAVIS Dataset [17]. The DAVIS Dataset has data on the interaction of 72 kinase inhibitors with 442 kinases covering >80% of the human catalytic protein kinome. The interaction indicator of DAVIS is Kd. Kd is the ratio of antibody dissociation rate (how fast it dissociates from antigen) to antibody binding rate (how fast it binds from antigen). Therefore, the smaller the Kd value, the greater the binding affinity. We can know the strength of the binding interaction between the drug and the target protein through the Kd value, that is, the strength of the binding affinity.

The dataset for binary classification was created as follows. There are 1,202,086 in BindingDB, and a total of 187,700 data were sampled when extracted as positive and negative according to the critical points mentioned in the [26] and [19] papers. In BindingDB, the ratio of positive and negative numbers was unbalanced (Positive: 162,085, Negative: 25,615). Random sampling was used to balance the positive and negative data, and the training data were constructed by randomly sampling 27,000 positives and 21,000 negatives. The valid data consisted of the remaining data, and 2,100 samples of negative and 2,300 samples of positive were randomly sampled. Excluding the training data and validation data from the original data, there are 132,785 positives and 2,515 negatives. In order to balance, 2,415 positives and 2,415 negatives were extracted. In addition, the test data is composed of a total of five data sets according to the “Korea Testing and Certification Institute” test regulations, so that abnormal performance such as overfitting can be checked. Data set validation and performance validation were performed under the supervision of the respective institutions.

Comparing the 5 test datasets of Test1-5, the negatives have the same data, while the positives have about 2,300 different data. Objective evaluation was conducted to find the optimal deep learning model through a total of 5 positive test datasets with a difference of 95%. Table 4 below shows the configuration of training, validation, and test datasets.

**Table 4.** Data used to train the model and its form.

| Dataset | Protein Num | Drug Num | Positive Num | Negative Num |
| --- | --- | --- | --- | --- |
| Total | 3,731 | 118,288 | 162,085 | 25,615 |
| Train | 2,739 | 39,298 | 27,000 | 21,000 |
| Valid | 1,131 | 4,258 | 2,300 | 2,100 |
| Test1 | 1,191 | 4,776 | 2,415 | 2,515 |
| Test2 | 1,185 | 4,767 | 2,415 | 2,515 |
| Test3 | 1,224 | 4,770 | 2,415 | 2,515 |
| Test4 | 1,180 | 4,767 | 2,415 | 2,515 |
| Test5 | 1,197 | 4,755 | 2,415 | 2,515 |

For the dataset for regression, the same dataset used in [20] was used for training and inference.

### Metrics

The evaluation index and loss function in our study are shown in Table 6 below.

**Table 5.**
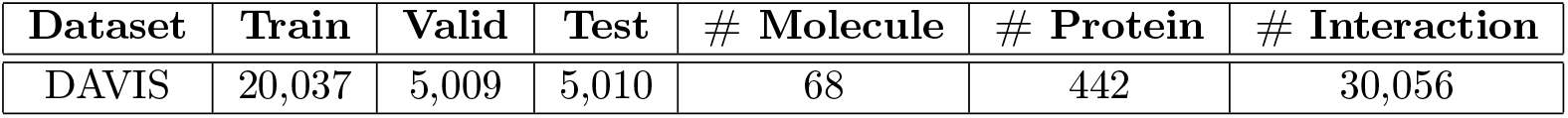
This is the dataset for regression.

**Table 6.** This is the cost function and Evaluation for Binary Classification and Regression.

|  | Binary | Regression |
| --- | --- | --- |
| Cost function | Binary Cross Entropy | Mean Square Error |
| Evaluation Metrics | Accuracy, AUPR | CI, AUPR |

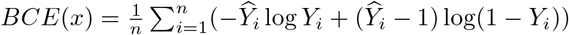

Where *Ŷ*_*i*_ is the *i*th label(1 for positive affinity and 0 for negative affinity) and *Y*_*i*_ is the *i*th predicted probability of being positive affinity for all *n* data. mMean Square Error

Where *Ŷ*_*i*_ is the *i*th label(Kd value : affinity score) and *Y*_*i*_ is the *i*th predicted affinity score for all *n* data.

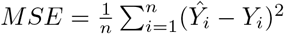

Where *ŷ* is followed by:

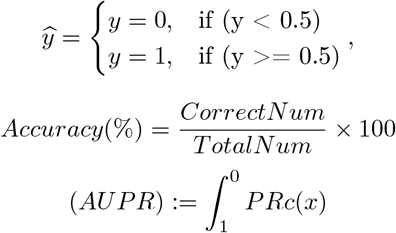

#### CI(Concordance Index)

Among the predicted affinity for the drug-target, two are randomly selected and whether the two are in the correct order is expressed as a probability value.

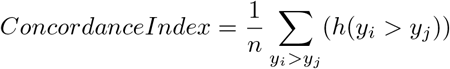

Where *h*(*x*) is followed by:

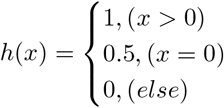

## Results

After the data preparation process, the experimental results for each model are shown in Tables 7 and 8 below. In the overall performance comparison, ABCnet performed the best, and in the case of regression, it did not outperform the SOTA model. There are three SOTA models used here. The first KronRLS is a similarity-based model that predicts DTI by combining a square error function with a specific regularization method [7]. The second SimBoost is also a similarity-based model based on a gradient boosting machine. Predict DTI through gradient descent using network metrics and PageRank scores [9]. Third, DeepDTA is a deep learning model that predicts DTI using drug and protein sequence pairs as input data. Fourth, GraphDTA is a graph deep learning model that predicts DTI using pairs of graphed data of drugs and protein sequences as input data. Concordance Index is similar to SOTA, but the score of MSE does not surpass.

**Table 7.** This is the Hyperparameters for Binary Classification and Regression.

| Item | Binary | Regression |
| --- | --- | --- |
| Epoch | 100 | 100 |
| Batch size | 512 | 256 |
| Optimizer | Adam | Adam |
| Learning rate | 0.001 | 0.001 |
| Protein Hidden layer Dimension | [128, 256, 512] | [128, 256, 512] |
| Drug Hidden layer Dimension | 512 | 512 |
| DTI Hidden layer Dimension | [256, 1] | [1024, 512, 1] |

**Table 8.** This is the result for Binary Classification.

| Dataset | MPNN | BCNet | ACNet | ABCnet |
| --- | --- | --- | --- | --- |
| Valid | 95.8409% | 95.85% | 95.9545% | 96.2727% |
| Test1 | 95.8603% | 95.4139% | 95.9212% | 96.6517% |
| Test2 | 95.9419% | 95.3129% | 95.8405% | 96.8144% |
| Test3 | 95.9347% | 95.2651% | 95.8874% | 96.7397% |
| Test4 | 95.8601% | 95.2767% | 95.7840% | 96.7885% |
| Test5 | 95.8806% | 95.3612% | 95.8238% | 96.7573% |

**Table 9.**
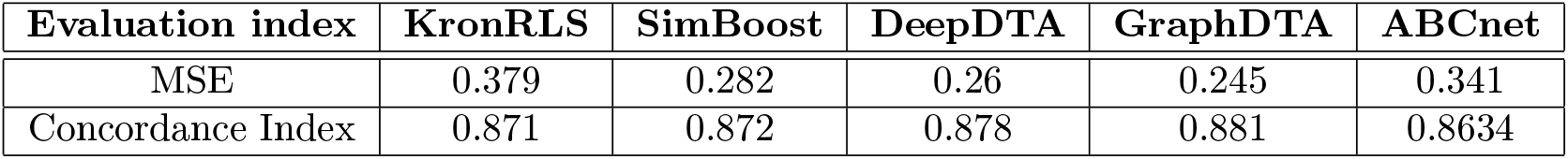
This is the result compare with SOTA.

https://www.overleaf.com/project/618a05533676801d8f68ccf6

## Discussion

ABCnet is a DTI deep-learning model that focuses on the drug model. Because we focused only on the drug model, the overall performance fell short of expectations. My goal is to focus on the drug model, taking one step towards finalizing an accurate DTI model in the future. The next thing to proceed is the study of the protein model. Looking at the deep learning research that predicts the current DTI, the input data used in the deep learning model uses a drug (molecule) sequence and a protein sequence. In particular, the protein sequence is a one-dimensional sequence of amino acids. However, when docking Molecule and Target, 3D material is docked in 3D space. Therefore, the current deep learning research that docks Molecule and Target as a one-dimensional sequence has a distance from the actual DTI. In order to narrow this distance, research to predict the 3D structure of Protein or further predict the 3D structure of Protein itself is absolutely necessary. Therefore, the next research that I will proceed is to complete the accurate protein model by including the 3D structure of the protein and further predicting it. It is not only to predict the simple binding value through the model that predicts the protein 3D structure, but also predicts the binding site to make a more accurate and useful model.

## Conclusions

We proposed an ABCNet model with Molecule Feature Embedding and Attention applied within the DTI model in the DTI model for predicting drug and target affinity. Among the affinity prediction problems, our model showed about 1% improvement in performance in comparison with SOTA in classification, and similar or lower performance in regression than SOTA. In the case of the regression problem, since the MSE is compared, it is not possible to know how good the model is because the difference in the size is so small, but it is meaningful in that the difference is not large. Combining these two results, we come to the conclusion that there was an improvement in performance compared to SOTA. In DTI research, like ours, most of the studies improve Molecule Feature Embedding. A study to strengthen the expression learning ability of Target Sequence Embedding is needed, and we plan to study this in the future.

## Acknowledgement

This work was supported by the Technology development Program(S2837441) funded by the Ministry of SMEs and Startups(MSS, Korea).

